# Composition and Abundance of Drifting Fish Eggs on the Upper Reaches of Xijiang River, China, after the Formation of the Cascade Reservoirs

**DOI:** 10.1101/2020.01.13.904110

**Authors:** Gao Minghui, Wu Zhiqiang, Tan Xichang, Huang Liangliang, Huang Haibo, Liu Hao

## Abstract

To develop effective management actions of riverine fisheries, it is important to monitor how fish resources (i.e., eggs) are recruited in the upper reaches of natural rivers, particularly where dams have been constructed, which potentially hinder life-history strategies. Here, we aimed to determine the State of drifting fish eggs resources, and the underlying environmental factors regulating the presence of fish eggs in the upper reaches of a river (Laibin section of Xijiang River, China). Based on surveys conducted over one spawning period (2016), we set out to: (1) describe the composition and abundance of drifting fish eggs in the 150 km Lainbin section under a dam control, and (2) analyze how the composition and distribution of fish eggs was correlated with environmental factors. A total of 15157 eggs belonging to two orders, four families, and 18 species were collected. Comparison of these data with historical records in the same area showed that the community structure of drifting eggs has changed considerably. Previously, the community was dominated by larger-bodied species, rather than the smaller species documented in 2016. Eggs were primarily detected between May and August. In the natural channel, the greatest abundance of eggs occurred during May and June. In comparison, the greatest abundance of eggs downstream of the dam was detected in July. The results of this study provide important information for water conservancy institutions towards managing regions containing dams to maintain the ecology of rivers and protect important fish resources.

## Introduction

The diversity of species globally is declining at an alarming rate because of human influences [1]. Global fishery resources are in decline due to many man-induced stressors, such as overfishing, dam construction, biological invasions, and climate change [2]. Aquatic ecosystems have been severely degraded as a result of anthropogenic changes to landscapes [3], with the damming of rivers representing a major anthropogenic factor impacting the ecology of freshwater fish populations. In particular, dams cause habitat loss, affect fish reproductive environments, and cut off migration routes [4]. Globally, 77% of rivers longer than 1000 km no longer flow freely from source to sea, due to obstructions by dams and reservoirs, with up- and downstream fragmentation and flow regulation being the leading contributors to the loss of river connectivity [5]. To develop practical and effective fishery management strategies, it is important to understand the recruitment of fish resources and monitor the status of breeding populations [6].

Since the 1980s, 216 fish species have been recorded in the Xijiang Basin in southern China, of which 30 species are endemic [7]. Various studies have shown that the spawning grounds of most fish species of economic importance are distributed in its middle and upper regions of this basin [8, 9]. However, the diversity and resources of fishes are under threat from dam construction. Already, the composition of fish species in the upper reaches of Xijiang River has been significantly impacted by the construction of a dam and the consequent creation of the Cascade Reservoirs [10].

Specifically, the Xijiang River Basin is rich in hydraulic resources. Eleven dams have been constructed on its upper and middle reaches, with the first being completed in 1980, and plans for the final dam (Datengxia) to be operational by 2020. This final structure will close off the last naturally flowing section in the upper Xijiang Basin. Historically, there were many spawning grounds for fish of economic importance in this section [8]; however, the spawning grounds of these species will be submerged once the dam is completed.

Most studies on fish reproductive ecology in Xijiang River have focused on larval fish in the middle and lower reaches [9, 11–13]. These studies have investigated annual dynamics in the abundance of fish larvae and their relationship with variation in hydrology, as well as patterns in their temporal distribution. However, few studies have examined the movement patterns of drifting fish eggs in the upper reaches of rivers [8]. Yet, it is important to monitor how fish resources are recruited in natural rivers, along with the status of breeding populations, to develop practical and effective fisheries management strategies [6, 14]. Studies of fish eggs provide information on ichthyology, environmental inventory, stock monitoring, and fisheries management. Patterns in the drift of fish eggs could provide insights on processes associated with spawning and larval production, as well as estimates of the stock size of spawning adults [15].

At present, one section of the upper reaches of Xijiang River remains free of dams (the Laibin section). Thus, here, we aimed to determine the underlying environmental factors regulating the presence of fish eggs in this section of the river. Specifically, we: (1) described the composition and abundance of drifting fish eggs in this section, and (2) analyzed how the composition and distribution of fish eggs was correlated with environmental factors. This information was gathered before this section of river was transformed into Datengxia Reservoir. The results of this study are expected to help inform water conservancy institutions on how to manage such regions to maintain the ecology and protect important fish resources.

## Materials & Methods

### Study area

Xijiang River is the largest tributary of the Pearl River, which is the largest river in south China. Xijiang River is 2214 km long, with a catchment of 353,120 km^2^ and mean annual discharge of 2.24 × 10^11^ m^3^. The upper reaches of Xijiang River are located in a geotectonically complex karst area, with mountains and valleys on either side. The river is composed of curved channels, beaches, underground streams, and karst caves [8]. The river is located in a region with a subtropical climate, with a mild climate and abundant rainfall. The river has high flow in summer and low flow in spring and winter. Since the 1980s, the landscape and the hydrodynamics of the basin have been modified by the construction of 11 hydroelectric power plants (HEPs). This study was carried out in the Laibin section of the river, which is under the influence of the Qiaogong and Honghua HEPs, spanning approximately 230 km of the upper reaches of Xijiang River (Fig. 1).

**Fig. 1.**
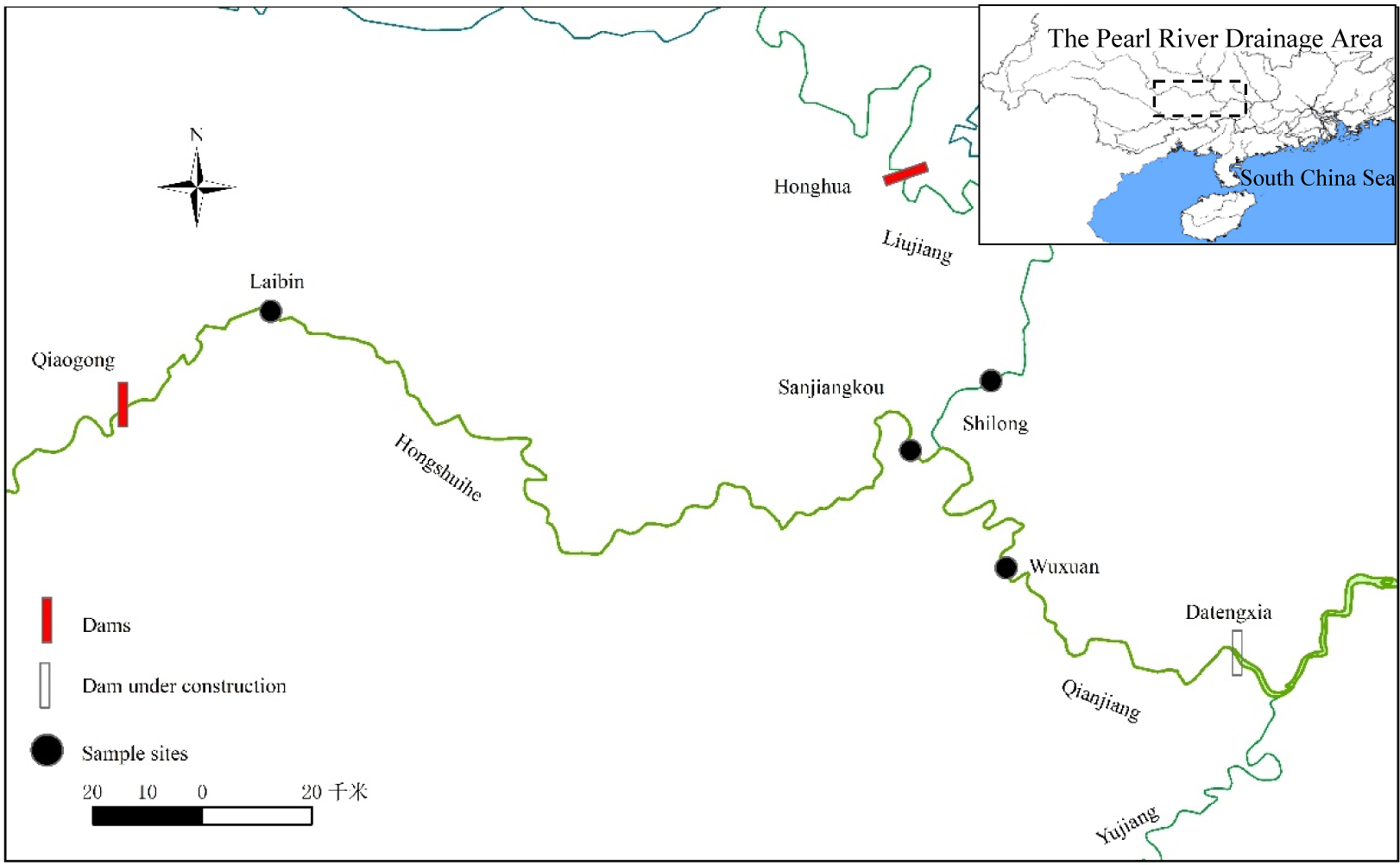
Upper reaches of the Xijiang River network, showing the sampling sites and dams.

### Sampling and data collection

Samples were collected daily during the spawning season (April to August) in 2016. Four sampling sites were set up to collect drifting fish eggs. These sites were located at Laibin (109° 14’ 31" E, 23° 42’ 41" N), Sangjiangkou (109° 32’ 6" E, 23° 47’ 36" N), Shilong (109° 32’ 10" E, 23° 50’ 29" N), and Wuxuan (109° 39’ 54" E, 23° 36’ 22" N). About 150 km separates Laibin from Wuxuan, with the four sampling sites being set at 60 km distances from one another. The four sites fell into three categories: (1) downstream of the dams (Laibin sites, which was downstream of the Qiaogong HEP); (2) at a junction of two rivers (Sangjiangkou and Shilong site, which were at the confluence the Hongshuihe and Liujiang Rivers); and (3) on a conventional channel (Wuxuan site, which was 55 km downstream of the confluence, and approximately 60 km upstream of the Datengxia HEP under construction).

Fish eggs samples were collected using Jiang nets (total length 5 m; rectangular iron opening/mouth 1.0 m × 1.5 m, and a mesh net size of 0.5 mm attached to a 0.8 m × 0.4 m × 0.4 m filter collection bucket). We selected sections with a width of 200 m and a depth of 12 m for sampling. The nets were deployed in the surface water, 10 m from the shore, to optimize the catch of target species. Sampling duration was 1 h (06:00–07:00), against the current.

Fish eggs were counted and sorted to the lowest possible taxonomic level at each station, based on morphological characteristics [16–18]. Eggs that could not be identified were placed in an aerated tank (diameter 30 cm, height 40 cm) at 20–25 °C for at least 1 week, until the species could be identified. Each tank was stocked with 40–50 eggs.

Real-time water temperature (WT) and dissolved oxygen (DO) data were collected using a dual input multiparameter digital analyzer (HACH HQ40d, Yiku Industrial Instrument Co., LTD, Shanghai China). Water transparency (Tra) was measured using a Secchi disk (HENGLING Technology Co., Ltd., Wuhan, China). Data on discharge (Dis), velocity (Vel), and water level (WL) were obtained from the website of the Pearl River Water Conservancy Commission (http://www.pearlwater.gov.cn.).

### Data analysis

The abundance of eggs was expressed as catch per unit effort (CPUE). The dominant species were determined using the Index of Relative Importance (IRI)[19]:

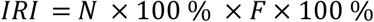

where N × 100 % and F × 100 % represent the relative abundance and frequency of occurrence, respectively. The IRI of the dominant species should exceed 100.

Spawning site location (*L*) was calculated as:

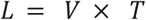

where T is the number of hours of development of the fish eggs when captured, based on their developmental stage (Table 1); and V is the water velocity at that moment (km/h).

To assess spatiotemporal variation in the distribution of eggs, a two-factor analysis of variance (ANOVA) was applied, in which differences were regarded as significant at 0.05 probability. Canonical correspondence analysis (CCA) is a robust method that directly correlates community data to environmental variables by constraining species ordination to a pattern that is correlated with environmental variables [22]. CCA was used to analyze the correlation between environmental factors and the temporal distribution of fish eggs. Only numerically abundant species were tested to avoid any spurious effects caused by groups of rare species. All maps were drawn using Surfer 8.0. Statistical analyses were performed using SPSS 20 (SPSS, IBM, Armonk, NY, USA) and CANOCO 4.5 (http://www.canoco5.com).

**Table 1.**
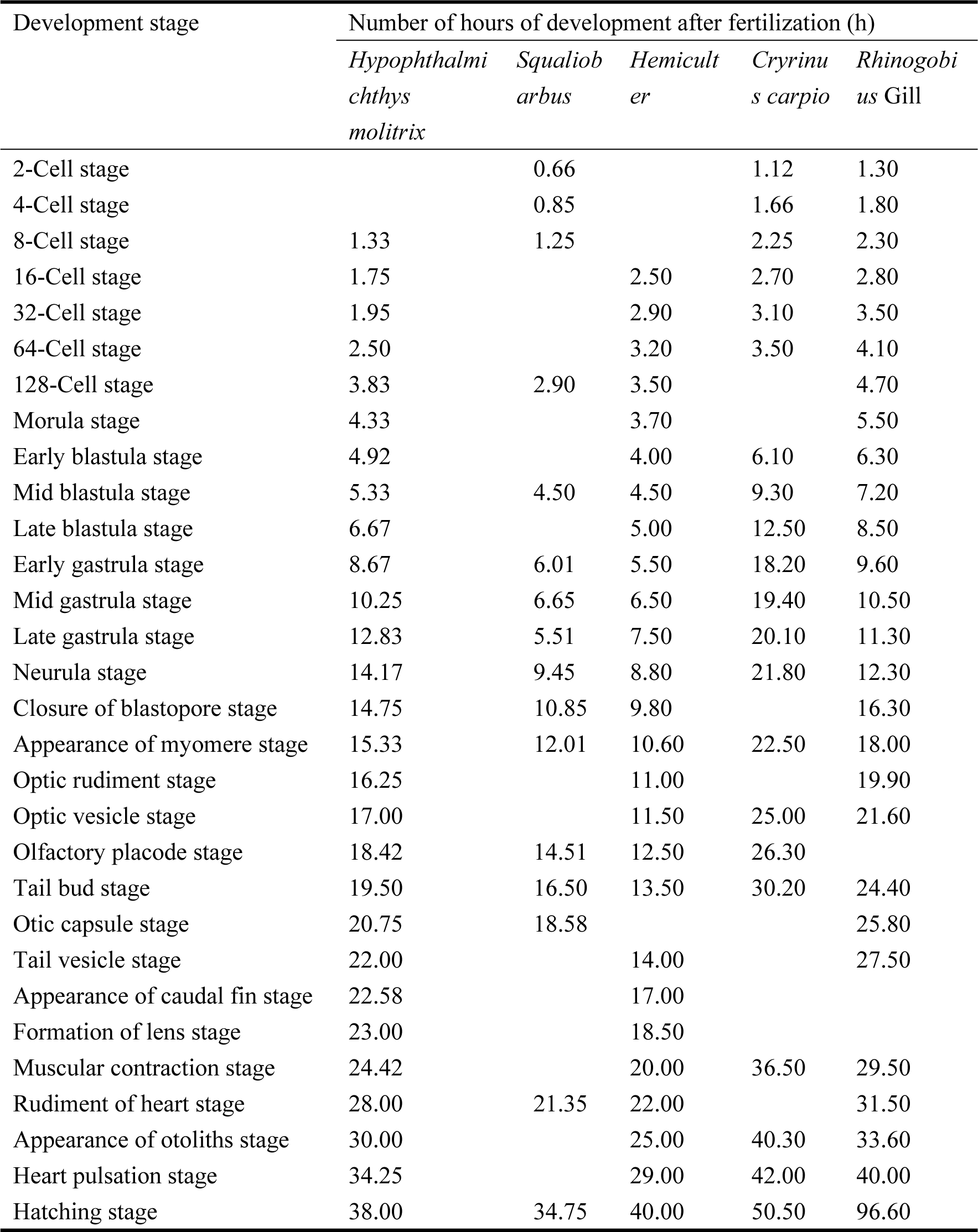
Number of development hours of eggs after fertilization for the different development stages of five of the sampled species (20–25 ℃) [6, 20, 21].

## Results

### Species composition of eggs

A total of 485 samples were collected during the surveys. Out of these samples, 15157 individuals were raised in aquaria to the larval or juvenile stage of fish. All specimens belonged to 18 species from four families and two orders, including three genera that could not be identified. The dominant species included *Squalidus argentatus* (30.67%), *Sinogastromyzon wui* (26.30%), and *Botia robusta* (10.50%). Other individuals belonged to the genera *Culter* Basilewsky, *Hemibarbu* Bleeker, and *Rhinogobius* Gill. Unidentified eggs represented 4.75% of all captured eggs (Table 2). Asian carp (*Hypophthalmichthys molitrix*) represented a small proportion of the collected samples. Only 22 eggs were found at the Shilong site (tributary sampling site) in July.

**Table 2.** List of species for which fish eggs were collected in the Laibin section of the river in 2016, showing the percentage of presence.

| Order | Family | Species | Station & Year |  |  |  | Total number of eggs | Ratio (%) | Development stages |
| --- | --- | --- | --- | --- | --- | --- | --- | --- | --- |
|  |  |  | LB | SJK | SL | WX |  |  |  |
| Cypriniformes | Cobitidae | *Botia robusta* | 36 | 1275 | 137 | 144 | 1592 | 10.50 | Tb - Hs |
|  |  |  | 4 | 417 | 562 | 48 | 1031 | 6.80 | Eg - Mc |
|  | Cyprinidae | *Squaliobarbus* | 1 | 420 | 445 | 65 | 931 | 6.14 | Tb - Hs |
|  |  | *curriculus* |  |  |  |  |  |  |  |
|  |  | *Pseudolaubuca* |  |  | 25 | 12 | 37 | 0.24 | Fl - Hs |
|  |  | *engraulis* |  |  |  |  |  |  |  |
|  |  | *Sinibrama macrops* |  | 88 | 23 | 18 | 129 | 0.85 | Tb - Hs |
|  |  | *Hemiculter* | 59 |  | 6 | 17 | 82 | 0.54 | Ns - Hp |
|  |  | *leucisculus* |  |  |  |  |  |  |  |
|  |  | *Culter Basilewsky* |  | 300 | 213 | 18 | 531 | 3.50 | Tb - Hs |
|  |  | *Hypophthalmichthys* |  |  | 22 |  | 22 | 0.15 | Mc |
|  |  | *molitrix* |  |  |  |  |  |  |  |
|  |  | *Hemibarbu Bleeker* |  | 136 |  | 12 | 148 | 0.97 | Fl - Hs |
|  |  | *Squalidus argentatus* | 30 | 2364 | 1937 | 318 | 4649 | 30.67 | Eb - Hp |
|  |  | *Saurogobio dabryi* | 33 | 242 | 276 | 75 | 626 | 4.13 | Ns - Hs |
|  |  | *Cirrhinus molitorella* |  | 10 | 43 | 52 | 105 | 0.69 | Tb - Hs |
|  |  | *Osteochilus salsburyi* |  | 391 | 139 | 47 | 577 | 3.80 | Ov - Rh |
|  |  | *Cryrinus carpio* |  | 29 | 69 | 13 | 111 | 0.73 | Mc - Hs |
|  |  | *Carassius auratus* |  | 9 | 49 | 31 | 89 | 0.58 | Mc - Hp |
|  | Homalopteridae | *Protomyzon sinensis* |  | 206 | 195 | 67 | 468 | 3.08 | Eg - Hp |
|  |  | *Sinogastromyzon wui* | 188 | 2631 | 832 | 335 | 3986 | 26.30 | Eg - Hp |
| Perciformes | Gobiidae | *Rhinogobius Gill* |  | 43 |  |  | 43 | 0.28 | Mc - Hs |
|  | Total |  | 351 | 8561 | 4973 | 1272 | 15157 | 100 |  |
|  | Number of species |  | 7 | 15 | 16 | 16 | 18 |  |  |

### Spatial and temporal variation

The abundance of eggs varied significantly across the three site categories and months of the study period (*P* < 0.05). The greatest relative abundance was observed at the junction of two rivers (Sangjiangkou and Shilong) in June 2016 (77 eggs/net/h). All identified species were detected at this site. The site with the second greatest abundance was the conventional channel (Wuxuan) in June 2016 (45 eggs/net/h), with 16 species being detected. The lowest abundance was detected downstream of Qiaogong HEP (Laibin) in July 2016 (7 eggs /net/h), with nine species being detected (Table 3 and Fig. 2).

**Table 3.**
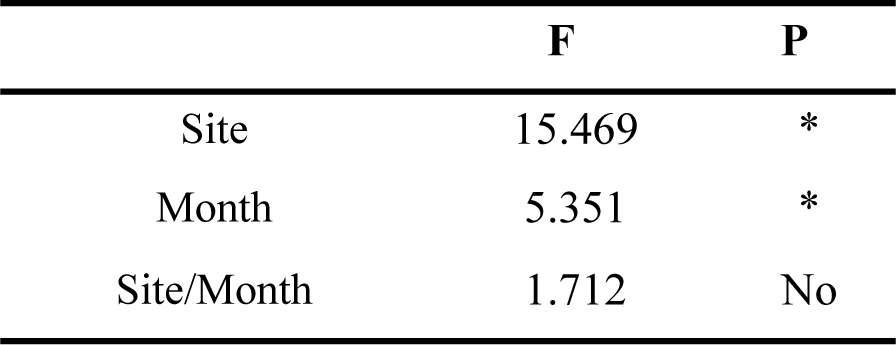
ANOVA on the spatio-temporal variation of fish egg distribution at the sampling sites. * Significantly different (p < 0.05)

|  | **F** | **P** |
| --- | --- | --- |
| Site | 15.469 | * |
| Month | 5.351 | * |
| Site/Month | 1.712 | No |

**Fig. 2.**
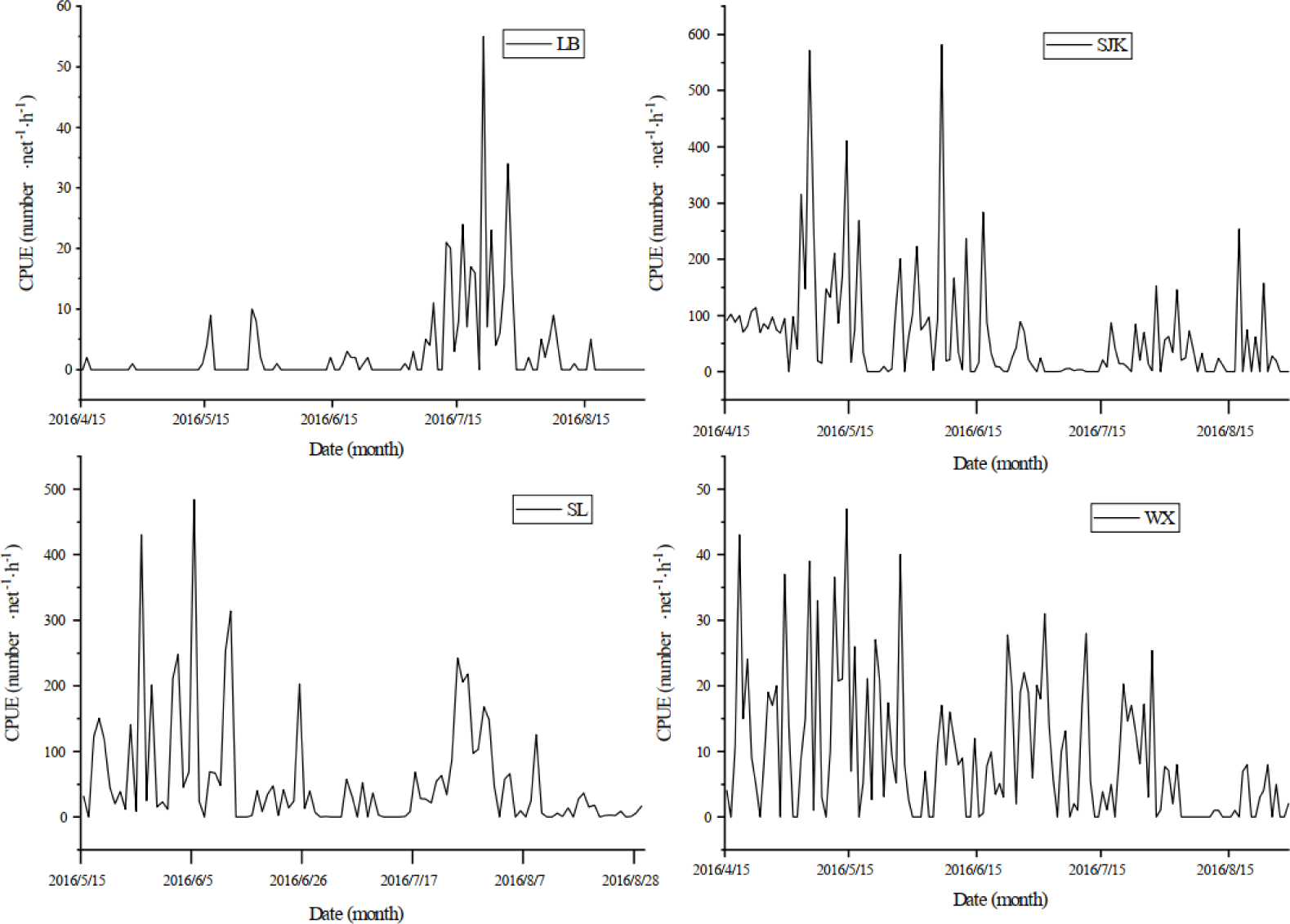
Monthly variation in the CPUE of drifting eggs at the sampling sites in 2016.

Drifting eggs were mainly found between May and July, with several spawning peaks occurring in each year (Fig. 2). The species composition of eggs differed across the sampling months (Fig. 3). The proportions of species belonging to Gobioninae increased from April to August, whereas those belonging to Botiinae and Homalopteridae decreased over the same period.

**Fig. 3.**
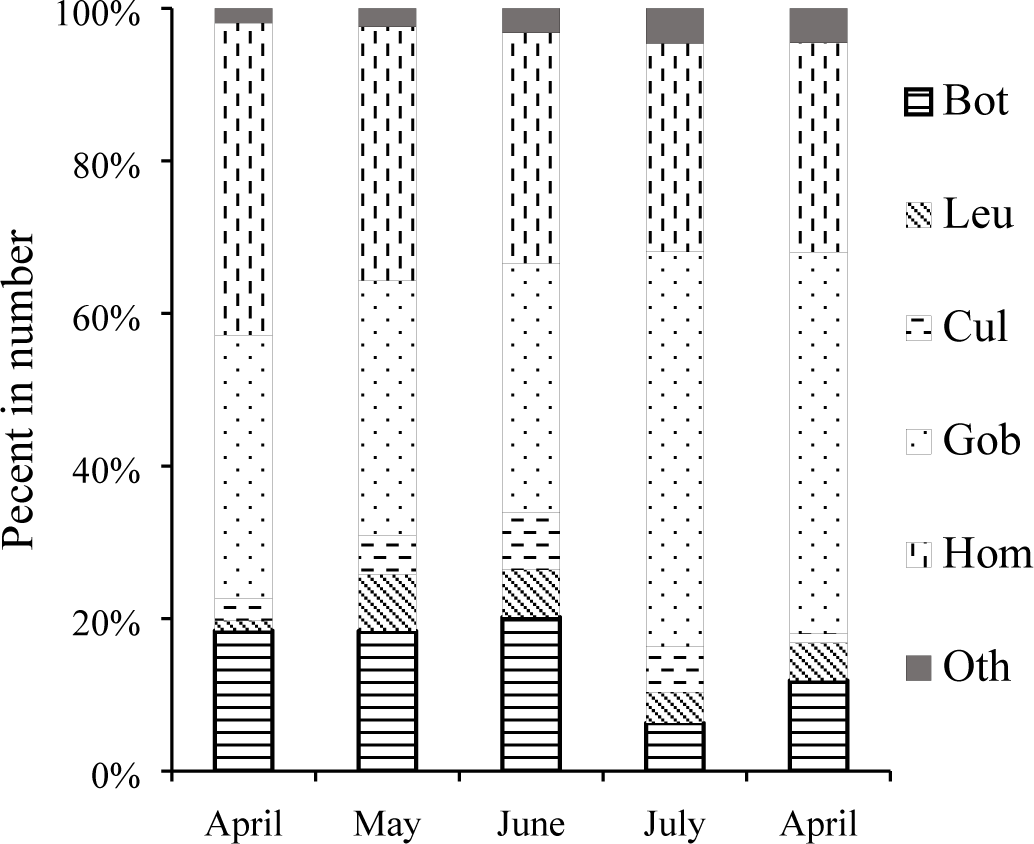
Percentage of eggs from the dominant groups of samples collected from the sampling sites from April to April of 2016. Bot, Botiinae; Leu, Lueciscinae; Cul, Cultrinae; Gob, Gobioninae; Hom, Homalopteridae; Oth, other.

### CCA analysis

CCA ordination was used to clarify the relationship between environmental factors and the abundance of eggs based on data from 13 species and a set of six environmental factors. Accumulated constrained eigenvalues for the first multivariate axes were 0.109 (CCA1) and 0.045 (CCA2), which denoted good species separation along the axis. The first four axes explained 92.85% of total variance. A Monte-Carlo test (Table 4) showed that WT and WL were the key environmental factors affecting assemblages (*P* < 0.05).

**Table 4.** Conditional effects and correlations of environmental variables based on the CCA axes

| Environmental | P | Axis 1 | Axis 2 |
| --- | --- | --- | --- |
| WT | 0.006 | 0.6294 | 0.5660 |
| WL | 0.002 | -0.0799 | 0.6009 |
| Vel | 0.066 | 0.4052 | -0.0583 |
| DO | 0.162 | -0.5284 | -0.1790 |
| Dis | 0.486 | -0.1228 | 0.5709 |
| Tra | 0.746 | 0.1218 | -0.5029 |

The CCA ordination plot of eggs (Fig. 4) showed that the correlation between environmental factors and the abundance of eggs varied with respect to species. *B. robusta* had a strong relationship with water level and dissolved oxygen. *Cirrhinus molitorella*, *Culter Basilewsky*, and *Saurogobio dabryi* were positively correlated with water level and temperature, but were less affected by dissolved oxygen and transparency. The presence of *Osteochilus salsburyi*, *Protomyzon sinensis*, and *Si. keangsiensi* was positively correlated with dissolved oxygen and transparency, but was less affected by water level and temperature.

**Fig. 4.**
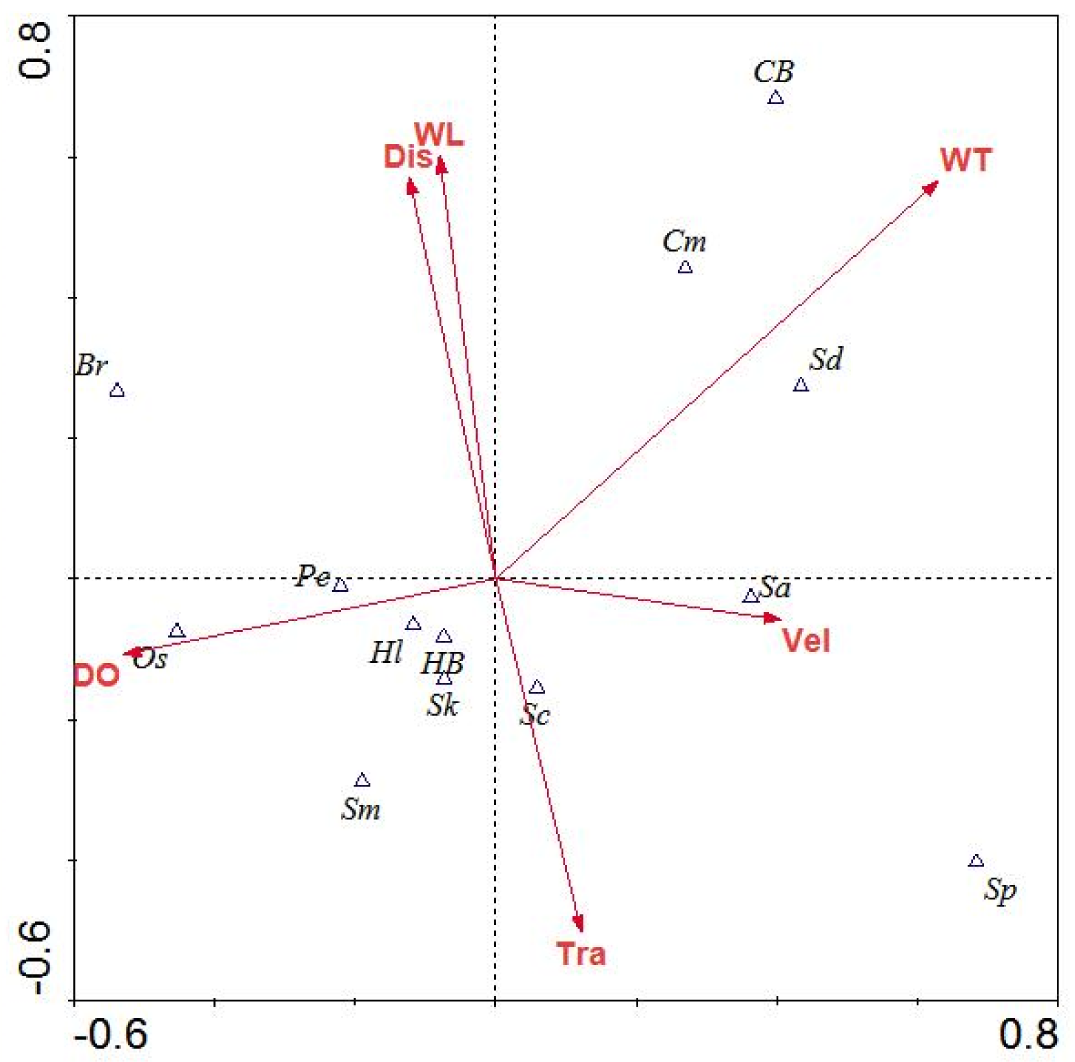
Canonical correspondence analysis showing how the eggs of different fish species are correlated with environmental variables on the two canonical axes. Vectors represent the correlation of environmental factors with the Canonical axes. Key to abbreviations: Br = *Botia robusta*, Sp = *Sinibotia pulchra*, Sc = S*iobarbus curriculus*, Pe = *Pseudolaubuca engraulis*, Sm = *Sinibrama macrops*, Hl = *Hemiculter leucisculus*, CB = *Culter* Basilewsky, HB = *Hemibarbu* Bleeker, Sa = *Squalidus argentatus*, Sd = *Saurogobio dabryi*, Cm = *Cirrhinus molitorella*, Os = *Osteochilus salsburyi*, Sk = *Sinohomaloptera keangsiensis*.

### Variation in spawning sites

The developmental stages of the 18 species captured in this study are presented in Table 2. Based on the data collected in this study, we calculated that the spawning sites of drifting fish eggs were 15–89 km upstream of our sampling sites. Three spawning grounds exist upstream of Laibin City, while five exist downstream, with the city being located 40 km downstream of Qiaogong HEP (Fig. 5). This finding differs to the forecasts of spawning grounds made before the Qiaogong HEP was completed [8]. Thus, the spawning grounds located upstream of Datengxia HEP will be completely submerged after impoundment. Our findings confirm the presence of spawning grounds in the Laibin section of Xijiang River.

**Fig. 5.**
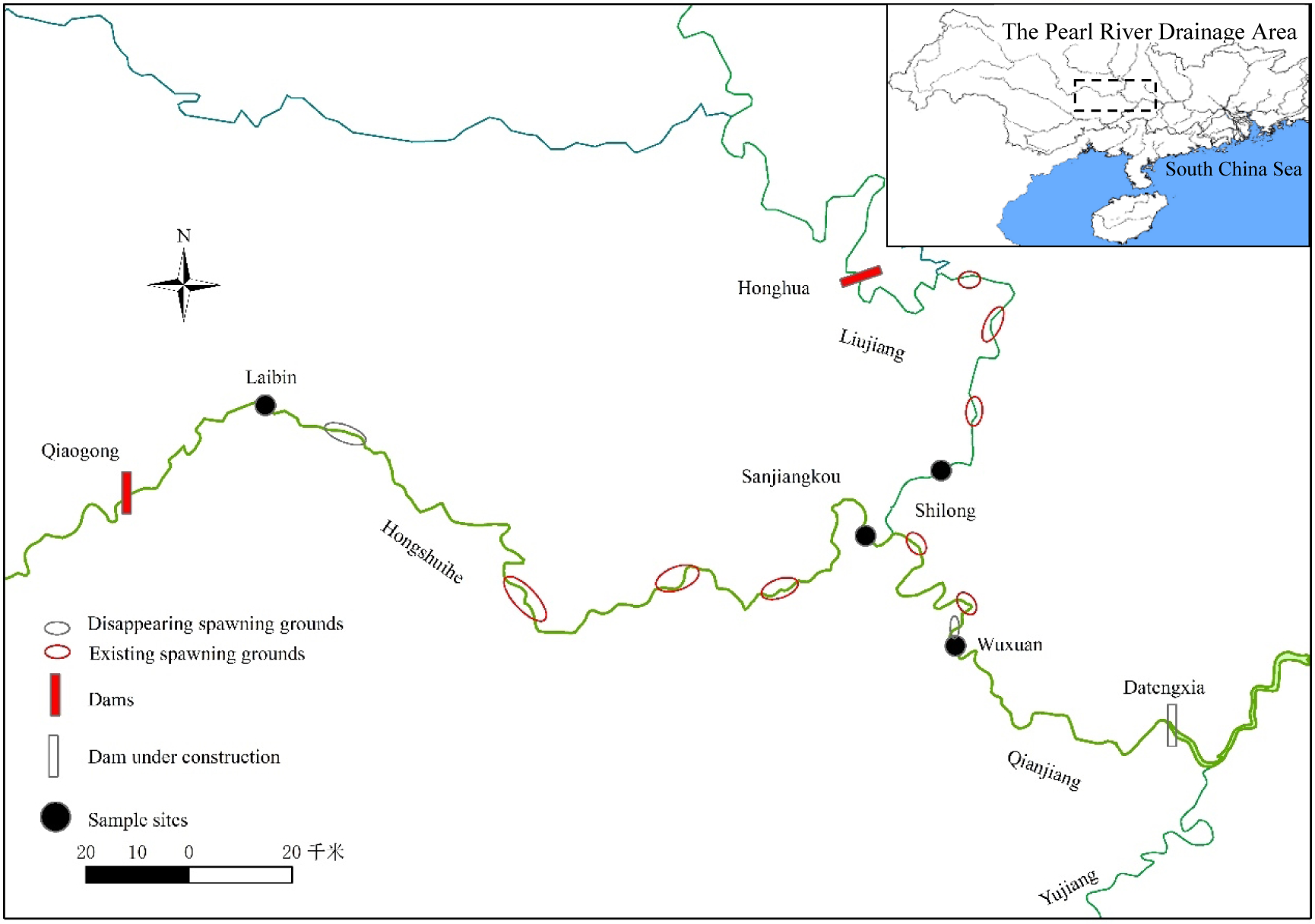
Distribution of spawning grounds in the upper Xijiang River.

## Discussion

### Variation in the community structure of drifting eggs

Knowledge remains limited about the composition and abundance of drifting fish eggs in the upper reaches of Xijiang River, southern China. This study demonstrated that assemblages of eggs in this region were mainly composed of species from the families Cobitidae, Cyprinidae, and Homalopteridae. This community has changed considerably compared to historical data [8]. The dominant species spawning in this region previously included *Mylopharyngodon piceus*, *Ctenopharyngodon idellus*, *Hy. molitrix*, *Hy. nobilis*, *Mystus guttatus*, *Squaliobarbus curriculus*, *Cirrhinus molitorella*, and *Cryrinus carpio*. In comparison, only *Sq. curriculus* was detected in the current study, representing 6.14% of all species, with all other dominant species having even smaller percentages. This shift in the structure of the fish community in the upper reaches of Xijiang River is likely due to anthropogenic changes to the hydrodynamics of the river, with the construction of the Cascade Reservoirs being the main driver [23]. In contrast to larger fish species, small-sized species tend to batch spawn small eggs over extended reproductive periods, in addition to parental care and migration being absent [24]. These characteristics might make these species better suited to the current environment of the upper Xijiang River following the development of the Cascade Reservoirs.

Asian carp was previously the most economically important taxon in this region (Zhou, 2005); however, we only found *Hy. molitrix* eggs at the Shilong sampling site in the Liujiang River during July 2016. The decline in Asian carp resources is a common phenomenon along the entire length of Xijiang River. The percentage of these species at the larval stage was 46.6% in 1986, but declined to 4.6% by 2008 [9]. These species have specific spawning requirements [25], including large, turbid rivers characterized by high turbulence caused by hard points (complex structure) or tributary confluences [26]. After the Cascade Reservoirs were constructed in the upper reaches of Xijiang River (2008), the flow regime changed markedly, with a decrease in water velocity and fewer periods with major floods (Wang, 2015). This change to the river might have contributed to the decline in spawning activity by Asian carp.

### Relationship between fish reproduction and environmental factors

Research remains limited on the reproductive patterns of fish in the upper reaches of Xijiang River. Yet, such information is essential for fisheries regulation and management to be effective. Our data showed that the abundance and distribution of drifting eggs of different species exhibited significant temporo-spatial differences. Fish reproduction is directly associated with various environmental factors, with water temperature representing key factor governing the spawning period of freshwater species [27]. Temperature strongly influences the dynamics of fish populations [28], by stimulating the gonads [29] and affecting spawning frequency [30]. Furthermore, flood pulses trigger spawning, especially for fish with drifting eggs [12, 31, 32]. Longer flood durations and greater flow discharge enhance fish production [33]. Our study demonstrated that water temperature was the key environmental factor affecting the assemblages of eggs in the upper reaches of Xijiang River (Table 4).

We confirmed the presence of reproduction at sampling sites located at a river junction and the absence of reproduction at sampling sites located downstream of dams. We also demonstrated that the reproductive peak occurred in May and June at the junction and conventional channel, but in July downstream of the dams. These constricted spawning periods and delayed spawning peaks might be caused by cold water present below dams (Tan, 2010). Hydrodynamic conditions at river junctions tend to be complex, particularly with respect to speed and the direction of currents. These parameters strongly influence the retention and dispersal of eggs. Our results support previous studies, which demonstrated that rheophilic fishes reproduce in such confluent areas [34].

The tributary Liujiang River is important in supplementing the fish resources of Xijiang River. Compared with the main stream, the Liujiang tributary contains a greater number of spawning species and has a longer spawning period, which could be explained by the greater abundance of multiple-spawning species, including *Sq. argentatus* and *Si. keangsiensis*. These species are generally classed as sedentary or short-range migratory species that have extended reproductive periods [35].

### Key threats to the reproduction and persistence of fishery resources

The current study showed that the dominant species that spawn drifting eggs were *Sinogastromyzon wui*, *Botia robusta*, *Sinibotia pulchra*, and *Squalidus argentatus* (Table 5). In particular, a significant correlation was detected between the reproduction of these species and water level. Water level was the main influencing factor in our study, and might facilitate the spawning of *Sinogastromyzon wui*, *Botia robusta* and *Sinibotia pulchra* in areas with cobblestones and sandbanks. With the planned operation of Datengxia Dam, the rise in water levels will cause these spawning grounds to become inundated. This phenomenon likely represents the most important potential threat to the spawning of species with floating fish egg resources in this section of the river.

**Table 5.**
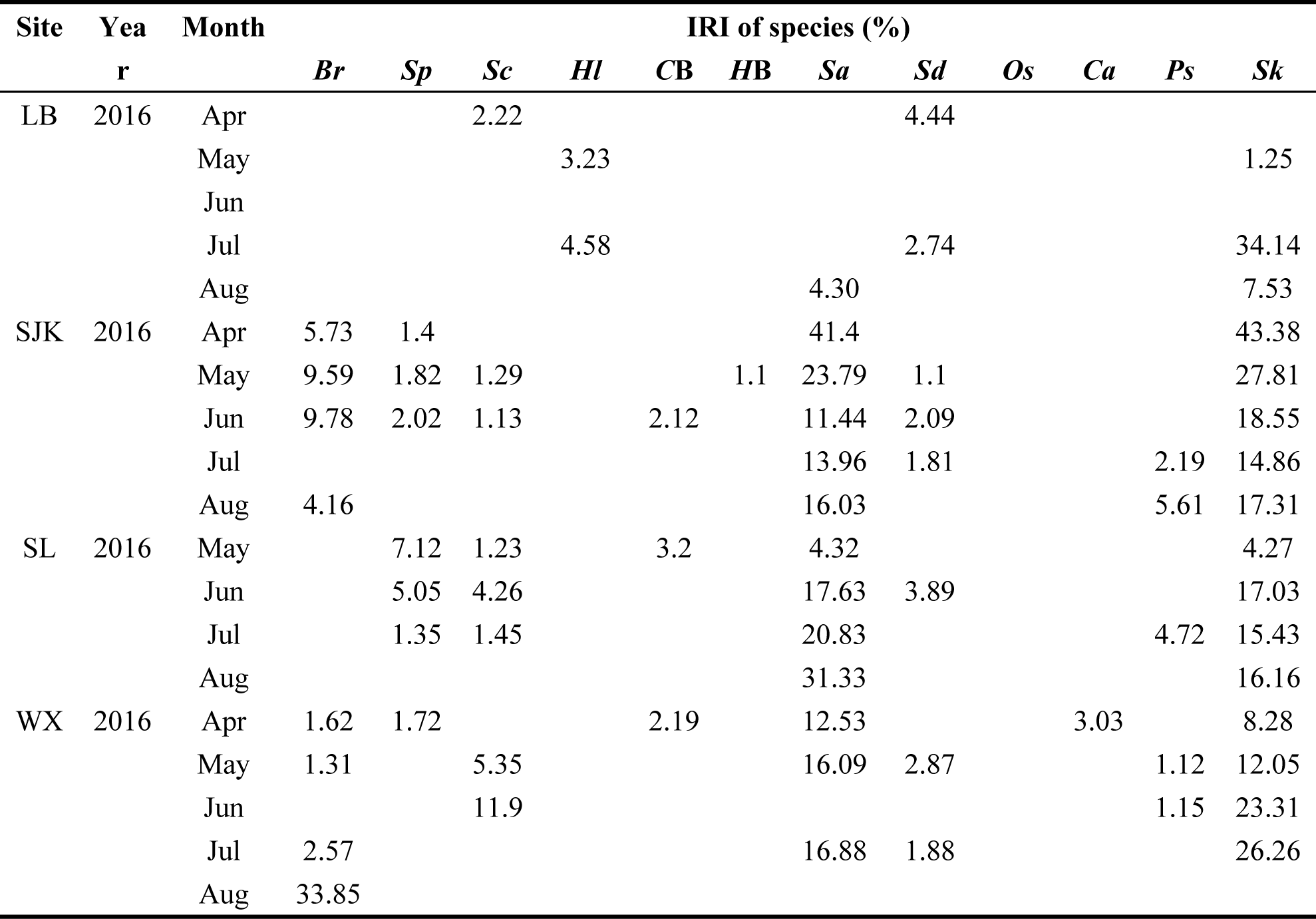
IRI of dominant species at the sampling sites.

The dominant species detected in the current study had extended reproductive seasons, spanning from April to August. At present, fishing is only banned from May until June in Xijiang River each year, failing to encompass the entire reproductive period. The reproductive period of fishes should be completely protected with no interference, especially for rare and threatened species. Conserving rare species by limiting the numbers caught implies that each individual in the population is valuable, with the removal of any individuals potentially causing a further decline in numbers [36]. In particular, the compensatory density dependence is assumed to be very weak in Yangtze River [6]. Consequently, the government has ordered a 10-year fishing ban in the Yangtze River from 2020. We suggest extending the fishing ban season to encompass the entire breeding period to be effective. In addition, the broad free stretches upstream of the reservoir in the tributary (Liujiang), will probably become important spawning grounds for many fishes following the onset of dam operation. For the effective conservation of fish recruitment, the fishery administration should strictly control fishermen’s nets and reduce the occurrence of electric fishing.

## Conclusions

In total, 15157 individuals of 18 species were collected over two surveys in 2016. We detected significant spatial and temporal variation in the number and abundance of the eggs of different species at the sampling sites. Water temperature and water level represented key environmental factors affecting the composition and abundance of eggs during the two survey periods, which contrasted with the results of previous studies. With rapid urbanization, freshwater fisheries and ichthyoplankton resources are noticeably declining. This phenomenon might further exacerbate the trend towards lower biodiversity in the upper reaches of Xijiang River. In conclusion, it is important to protect and continue monitoring fishery resources in Xijiang River to understand and mitigate the impacts of dam operations.

## Acknowledgements

We thank all the editors and reviewers for providing constructive comments on the present work.

## Additional information and declarations

### Author Contributions

All authors conceived and designed the experiments; MG performed the experiments; MG analyzed the data; MG and ZW contributed reagents/materials/analysis tools; MG prepared figures and/or tables; all authors wrote and reviewed drafts of the manuscript, and approved the final version.

- Minghui Gao conceived and designed the experiments, performed the experiments, analyzed the data, contributed reagents/materials/analysis tools, prepared figures and/or tables, authored and reviewed drafts of the paper, approved the final draft.
- Zhiqiang Wu conceived and designed the experiments, contributed reagents/materials/-analysis tools, authored and reviewed drafts of the paper, approved the final draft.
- Xichang Tan conceived and designed the experiments, authored and reviewed drafts of the paper, approved the final draft.
- Liangliang Huang conceived and designed the experiments, authored and reviewed drafts of the paper, approved the final draft.

